# Towards standardization of immune functional assays

**DOI:** 10.1101/718627

**Authors:** William Mouton, Chloé Albert Vega, Mathilde Boccard, François Bartolo, Guy Oriol, Jonathan Lopez, Alexandre Pachot, Julien Textoris, François Mallet, Karen Brengel-Pesce, Sophie Trouillet-Assant

**Affiliations:** Joint Research Unit Hospices Civils de Lyon-bioMérieux, Hospices Civils de Lyon, Lyon Sud Hospital, Pierre-Bénite, France; Medical Diagnostic Discovery Department (MD3), bioMérieux S.A., Marcy l’Etoile, France; Virologie et Pathologie Humaine - Virpath Team, Centre International de Recherche en Infectiologie (CIRI), INSERM U1111, CNRS UMR5308, ENS Lyon, Claude Bernard Lyon 1 University, Lyon, France; Soladis, Lyon, France; Hospices Civils de Lyon, Plateforme de Recherche de Transfert en Oncologie, Department of Biochemistry and Molecular biology, Lyon Sud Hospital, Pierre-Bénite, France; Joint Research Unit Hospices Civils de Lyon-bioMérieux, EA 7426 Pathophysiology of Injury-Induced Immunosuppression, PI3, Claude Bernard Lyon 1 University, Edouard Herriot Hospital, Lyon, France; Hospices Civils de Lyon, Department of Anesthesia and Critical Care Medicine, Edouard Herriot Hospital, Lyon, France; Hospices Civils de Lyon, Immunology Laboratory, Edouard Herriot Hospital, Lyon, France; Université de Lyon, Claude Bernard Lyon 1 University, Faculté de Médecine Lyon Est, Lyon, France; Centre de Recherche en Cancérologie de Lyon, INSERM U1052, CNRS UMR5286, Lyon, France

**Keywords:** Immune Functional Assay, Host Response, Stimulation, Independent Validation, Standardization

## Abstract

Recent advances in the immunotherapy field require evaluation of the immune function to adapt therapeutic decisions. Immune functional assays (IFA) are able to reveal the immune status and would be useful to further adapt/improve patient’s care. However, standardized methods are needed to implement IFA in clinical settings. We carried out an independent validation of a published method used to characterize the underlying host response to infectious conditions using an IFA. We evaluate the reproducibility and robustness of this IFA and associated readout using an independent healthy volunteers (HV) cohort. Expression of a 44 genes-signatures and IFNγ protein secretion and gene-expression was assessed after stimulation. We observed a strong host-response correlation between the two cohorts. We also highlight that standardized methods for immune function evaluation exist and could be implemented in larger-scale studies. This IFA could be a relevant tool to reveal innate/adaptive immune dysfunction in immune-related disorders patients.

## 1. Introduction

The increase in patients suffering from immune-related disorders led to the improvement in precise knowledge concerning the components and function of the immune system. Whether of physiological or pathophysiological origin, *i.e.* immunosenescence [1] or autoimmune diseases [2, 3], the identification of immune alterations requires close monitoring of the immune status. Their ever-increasing number and severity make these disorders a major public health problem. Several therapies targeting the immune system, such as anti-inflammatory agents or immune stimulatory therapy, are currently used to re-balance the immune system in immunocompromised patients. Nevertheless, clinicians lack the tools to decipher immune status, and more precisely, immune function, in order to adapt therapeutic decisions and personalize patient care.

In some clinical situations, the use of immune functional assays (IFAs) could help improve management and reveal valuable information compared to the use of circulating biomarkers such as cell counting or measurement of soluble and cell surface biomarkers. Based on the evaluation of the immune response after an *in vitro* stimulation with antigen, IFAs allow assessing the immune function as well as anticipate the ability to respond to therapy or fight a pathogen. In the last few years, T-cell mediated immunity (blood-based) assays, such as interferon gamma release assays (IGRA), have been developed and have revolutionized the detection of latent tuberculosis infection (LTI). This type of test enables clinicians to be more accurate and effective in the management of patients suffering from immune-related disorders. Research, as well as clinical studies, use and adapt these IFAs to try to respond to specific questions, such as the capacity of specific patients to respond to vaccination [4 – 6]. Despite encouraging results, the main challenge complicating the implementation of IFAs in clinical settings is the need for standardization in order to perform these methods in any laboratory, in a reliable and reproducible way [7].

Such tests should minimize the number of steps during sample preparation and ideally use the biological matrix directly sampled from the patient to limit technical variability and eliminate the pre-analytical error. As already described for cytokine release assay [8,9], use of direct measurement in whole blood sampling would be the relevant solution to minimize the risk of contamination and avoid causing non-specific cell activation or cell death [10]. Indeed, the use of whole blood rather than peripheral blood mononuclear cell (PBMC) isolation preserves all interactions between circulating immune cells and reflects the internal environment of the subject [11]. Nevertheless, in addition to this pressing need for standardization, the second difficulty encountered is the interindividual variability exhibited by the immune system.

In this context, the “Milieu Intérieur” team (MI) used a whole blood, semi-closed stimulation system built into syringe-based medical devices, with the ambition of defining the boundaries of a healthy immune response [12]. Using a cohort of 25 healthy volunteers (HV) (aged between 30–39 years) stratified by gender (13 women, 12 men), they challenged whole blood with complex stimuli (mimicking fungi, bacteria, and virus infection) and established a 44-gene expression signature, capturing the diversity of complex immune responses. In this study, independent validation of the protocol developed by Urrutia *et al.* was performed in order to evaluate the reproducibility of the technical process before its potential implementation in clinical settings.

## 2. Material and methods

### 2.1 Blood stimulation

According to EFS standardized procedures for blood donation and to provisions of the articles R.1243–49 and following ones of the French public health code, a written non-opposition to the use of donated blood for research purposes was obtained from HV. The blood donors’ personal data were anonymized before transfer to our research laboratory. We obtained the favourable notice of the local ethical committee (Comité de Protection des Personnes Sud-Est II, Bâtiment Pinel, 59 Boulevard Pinel, 69,500 Bron) and the acceptance of the French ministry of research (Ministère de l’Enseignement supérieur, de la Recherche et de l’Innovation, DC-2008-64) for handling and conservation of these samples.

Whole blood from healthy volunteers (n=20), stratified by gender (10 women and 10 men), and with a median age of 52 years (range: 25-65 years), was distributed (1 mL) into pre-warmed TruCulture tubes (Myriad Rbm, Austin, TX, USA) containing either medium alone (Null), staphylococcal enterotoxin B (SEB; 400 ng/mL), or lipopolysaccharide (LPS; 100 ng/mL (E.coli, O55:B5)). These were then inserted into a dry block incubator and maintained at 37°C. After 24 hours of stimulation, the pellet was separated from the supernatant using a valve and kept in TRI Reagent® LS (Sigma-Aldrich, Saint-Louis, MO, USA).

### 2.2 Cytokine and transcriptomic assays

Interferon gamma (IFN-*γ*) release was quantified from Truculture’s supernatant using an immunoassay in the VIDAS^®^ platform (bioMérieux, Marcy l’Etoile, France). As previously described, RNA extraction was performed using a modified protocol of the NucleoSpin 96 RNA tissue kit (Macherey-Nagel Gmbh&Co. KG, Düren, Germany) and using a vacuum system [12]. Nanostring technology was used for mRNA detection of a 44 gene panel (details in supplementary table S1), an hybridization-based multiplex assay characterised by its amplification-free step; 300 ng of RNA were hybridized to the probes at 67°C for 18 hours using a thermocycler (Biometra, Tprofesssional TRIO, Analytik Jena AG, Jena, Germany). After removal of excessive probes, samples were loaded into the nCounter Prep Station (NanoString Technologies, Seattle, WA, USA) for purification and immobilization onto the internal surface of a sample cartridge for 2-3 hours. The sample cartridge was then transferred and imaged on the nCounter Digital Analyzer (NanoString Technologies) where colour codes were counted and tabulated for the 44 genes.

### 2.3 Genes expression analysis

As previously described, each sample was analyzed in a separate multiplexed reaction that each included eight negative probes and six serial concentrations of positive control probes. Negative control analysis was performed to determine the background for each sample. Data was imported into nSolver^®^ analysis software (version 4.0) for quality checking and normalization of data (details in supplementary material and method S1) [12].

## 3. Results

### 3.1 Transcriptomic response

Immune response after stimulation with SEB (Fig. 1A) or LPS (Fig. 1C) showed a strong correlation between the two independent cohorts for the 44-gene expression signature, confirmed by Deming regression on medians (Spearman’s correlation coefficient rho= 0.99 for SEB and 0.90 for LPS; Fig. 1B and 1D) and on inter-deciles (Spearman’s correlation coefficient rho=0,96 for SEB and 0,94 for LPS; details in supplementary figure S1).

**Figure 1.**
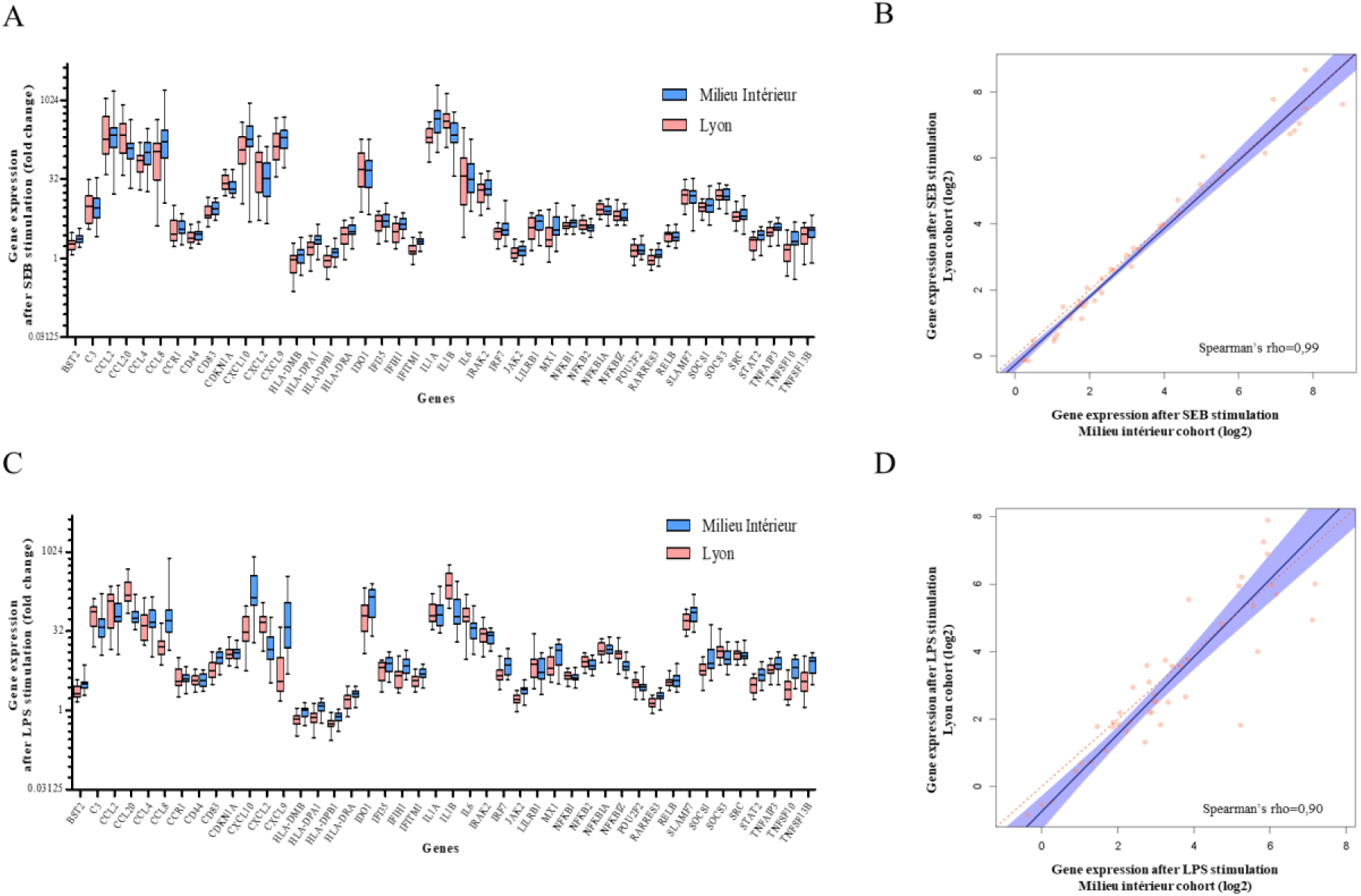
Gene expression after SEB- and LPS-whole blood stimulation. Whole blood from 25 healthy volunteers enrolled in the Milieu Intérieur cohort (blue) and from 20 healthy volunteers included in the Lyon study cohort (red) was stimulated in TruCulture tubes during 24h. After mRNA extraction, gene expression was quantified using Nanostring nCounter^®^ technology and normalized using NSolver^®^ software using four housekeeping genes (*HPRT1, POLR2A, RPL19* and *TBP*). Fold changes are used to describe gene expression variation between the stimulated condition (SEB (A) or LPS (C)) and the control condition (NULL) for the 44-gene signature studied. The Milieu Intérieur data was obtained from Gene Expression Omnibus (GEO) DataSets (GSE85176) [12]. Comparison of both cohorts was performed using a Deming regression analysis on gene expression medians following SEB (B) or LPS (D) stimulation. A Spearman’s correlation coefficient was calculated for each condition and indicated on each graphical representation. The orange dots represent individual gene expression, the blue beam represents the confidence interval, the blue line the Deming regression and the red dotted line the theoretical 100% identity.

### 3.2 Interferon gamma response

In a second time, following SEB stimulation, a comparison of *ifn-γ* gene expression (in the corresponding TruCulture) and IFN-*γ* release in the supernatant (Fig. 2) was performed. The latter is described as a marker of T-cell mediated immunity and used as a diagnostic tool in research and clinical settings (IGRA) [13,14]. A strong correlation was observed between protein and gene expressions for the 20 HV from Lyon (Fig. 2A) and the 25 HV from MI (Fig. 2B) (Spearman’s correlation coefficient rho ≥0.64 for both cohorts).

**Figure 2.**
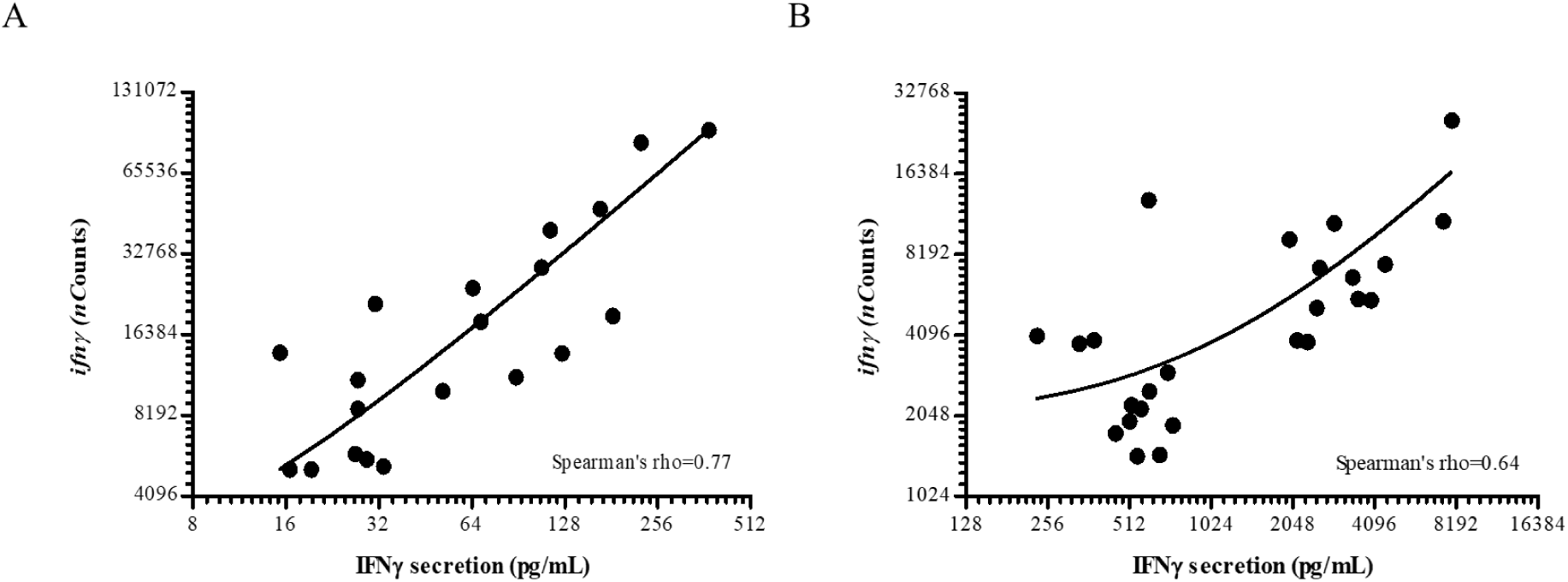
Interferon-*γ* induction after whole-blood SEB-stimulation. Whole blood from 20 healthy volunteers included in the Lyon study cohort (A) and from 25 healthy volunteers enrolled in the Milieu Intérieur (MI) cohort (B) was stimulated with SEB in TruCulture tubes for 24h. Comparison between IFNy mRNA and protein expression was assessed separately for each healthy cohort due to the use of a different method for cytokine secretion measurement. For protein quantification, IFN*γ* release was quantified in TruCulture supernatant using Luminex xMAP^®^ for the MI cohort and using VIDAS^®^ platform (bioMérieux) for the Lyon cohort. Count number for mRNA *ifn-γ*-gene expression, were normalized by the geometric mean of *HPRT1, POLR2A, RPL19* and *TBP* housekeeping genes. Gene data from the MI study was obtained from Gene Expression Omnibus (GEO) DataSets (GSE85176) [12] and protein data extracted from *Duffy et.al* supplementary information [15]. Spearman’s correlation coefficient (rho) was calculated for each condition and indicated on each graphical representation.

## 4. Discussion and conclusion

In this study, we carried out an independent validation of a published method used to characterize the underlying host response to infectious conditions using an IFA. We evaluate the reproducibility and robustness of this IFA and associated readout using an independent healthy volunteers (HV) cohort.

The strong host-response correlation between the two cohorts highlights that standardized methods for immune function evaluation exist and could be implemented in larger-scale studies. Also, based on the strong correlation observed, IFN-*γ* mRNA measurement might be a reliable and robust readout to assess T-cell functionality. Besides, the 24-hour stimulation of whole blood in standardized TruCulture tubes, which limits technical manipulation bias, renders this method a versatile tool for clinical use [11].

In conclusion, this standardized IFA could be a relevant tool to reveal innate and adaptive immune dysfunction in patients suffering from immune-related disorders.

## Supporting information

Supplementary informations

## Acknowledgements

We thank Pauline Desormeaux and Isabelle Mosnier for their technical assistance concerning nanostring molecular biology. We thank Verena Landel for language editing and critical reading of the manuscript.

## Conflict of Interest Disclosure

William Mouton, Chloé Albert Vega, Guy Oriol, Alexandre Pachot, Julien Textoris, François Mallet and Karen Brengel-Pesce, are employed at bioMérieux, an *in-vitro* diagnostic company, François Bartolo is an employee of Soladis, a services company specialized in data-related projects management,

## References

[1] J.J. Goronzy, C.M. Weyand, Understanding immunosenescence to improve responses to vaccines, Nat. Immunol. 14 (2013) 428–436. doi:10.1038/ni.2588.

[2] D.A. Smith, D.R. Germolec, Introduction to immunology and autoimmunity., Environ. Health Perspect. 107 (1999) 661–665. doi:10.1289/ehp.99107s5661.

[3] Y. Zhang, J.D. Rowley, Leukemias, Lymphomas, and Other Related Disorders, in: Emery Rimoins Princ. Pract. Med. Genet., Elsevier, 2013: pp. 1–44. doi:10.1016/B978-0-12-383834-6.00079-3.

[4] I. Macchia, F. Urbani, E. Proietti, Immune Monitoring in Cancer Vaccine Clinical Trials: Critical Issues of Functional Flow Cytometry-Based Assays, BioMed Res. Int. 2013 (2013) 1–11. doi:10.1155/2013/726239.

[5] T.M. Clay, A.C. Hobeika, P.J. Mosca, H.K. Lyerly, M.A. Morse, Assays for monitoring cellular immune responses to active immunotherapy of cancer, Clin. Cancer Res. Off. J. Am. Assoc. Cancer Res. 7 (2001) 1127–1135.

[6] A. Conrad, M. Boccard, F. Valour, V. Alcazer, A.-T. Tovar Sanchez, C. Chidiac, F. Laurent, P. Vanhems, G. Salles, K. Brengel-Pesce, B. Meunier, S. Trouillet-Assant, F. Ader, VaccHemInf project: protocol for a prospective cohort study of efficacy, safety and characterisation of immune functional response to vaccinations in haematopoietic stem cell transplant recipients, BMJ Open. 9 (2019) e026093. doi:10.1136/bmjopen-2018-026093.

[7] C. Albert-Vega, D.M. Tawfik, S. Trouillet-Assant, L. Vachot, F. Mallet, J. Textoris, Immune Functional Assays, From Custom to Standardized Tests for Precision Medicine, Front. Immunol. 9 (2018) 2367. doi:10.3389/fimmu.2018.02367.

[8] C.W. Thurm, J.F. Halsey, Measurement of Cytokine Production Using Whole Blood, in: J.E. Coligan, B.E. Bierer, D.H. Margulies, E.M. Shevach, W. Strober (Eds.), Curr. Protoc. Immunol., John Wiley & Sons, Inc., Hoboken, NJ, USA, 2005: p. im0718bs66. doi:10.1002/0471142735.im0718bs66.

[9] B. Yang, T.-H. Pham, R. Goldbach-Mansky, M. Gadina, Accurate and Simple Measurement of the Pro-inflammatory Cytokine IL-1β using a Whole Blood Stimulation Assay, J. Vis. Exp. (2011) 2662. doi:10.3791/2662.

[10] P. Brodin, D. Duffy, L. Quintana-Murci, A Call for Blood—In Human Immunology, Immunity. 50 (2019) 1335–1336. doi:10.1016/j.immuni.2019.05.012.

[11] D. Duffy, V. Rouilly, C. Braudeau, V. Corbière, R. Djebali, M.-N. Ungeheuer, R. Josien, S.T. LaBrie, O. Lantz, D. Louis, E. Martinez-Caceres, F. Mascart, J.G. Ruiz de Morales, Ottone L. Redjah, N.S.-L. Guen, A. Savenay, M. Schmolz, A. Toubert, M.L. Albert, Standardized whole blood stimulation improves immunomonitoring of induced immune responses in multi-center study, Clin. Immunol. 183 (2017) 325–335. doi:10.1016/j.clim.2017.09.019.

[12] A. Urrutia, D. Duffy, V. Rouilly, C. Posseme, R. Djebali, G. Illanes, V. Libri, B. Albaud, Gentien B. Piasecka, M. Hasan, M. Fontes, L. Quintana-Murci, M.L. Albert, L. Abel, A. Alcover, K. Astrom, P. Bousso, P. Bruhns, A. Cumano, C. Demangel, L. Deriano, J. Di Santo, F. Dromer, G. Eberl, J. Enninga, J. Fellay, A. Freitas, O. Gelpi, I. Gomperts-Boneca, S. Hercberg, O. Lantz, C. Leclerc, H. Mouquet, S. Pellegrini, S. Pol, L. Rogge, A. Sakuntabhai, O. Schwartz, B. Schwikowski, S. Shorte, V. Soumelis, F. Tangy, E. Tartour, A. Toubert, M.-N. Ungeheuer, L. Quintana-Murci, M.L. Albert, Standardized Whole-Blood Transcriptional Profiling Enables the Deconvolution of Complex Induced Immune Responses, Cell Rep. 16 (2016) 2777–2791. doi:10.1016/j.celrep.2016.08.011.

[13] Q. Yin, S. El-Ashram, H. Liu, X. Sun, X. Zhao, X. Liu, X. Suo, Interferon-Gamma Release Assay: An Effective Tool to Detect Early Toxoplasma gondii Infection in Mice, PLOS ONE. 10 (2015) e0137808. doi:10.1371/journal.pone.0137808.

[14] M. Ang, W. Wong, C.C.L. Ngan, S.-P. Chee, Interferon-gamma release assay as a diagnostic test for tuberculosis-associated uveitis, Eye. 26 (2012) 658–665. doi:10.1038/eye.2012.1.

