## Supplementary informations for "Towards standardization of immune functional assays"

Supplementary material and method S1

*Genes expression analysis*

A first step of normalization using the internal positive controls allowed for correction of potential sources of variation associated with the technical platform. To do so, the geometric mean of the positive probe counts was calculated for each sample. A mean of geometric means was then calculated for all patients. A scaling factor for each patient was determined as follows: geometric mean of the patient / mean of geometric means for all patients. For each patient, each gene count was divided by the corresponding scaling factor. Next, the background level was calculated as the mean +2 SD across the six negative probe counts. Finally, the method used for positive control normalization was applied to normalize for differences in RNA input, except that geometric means were calculated over four housekeeping genes (*RPL19, TBP, POLR2A*, and *HPRT1*) according to the Milieu Intérieur study [12].

Supplementary figure S1

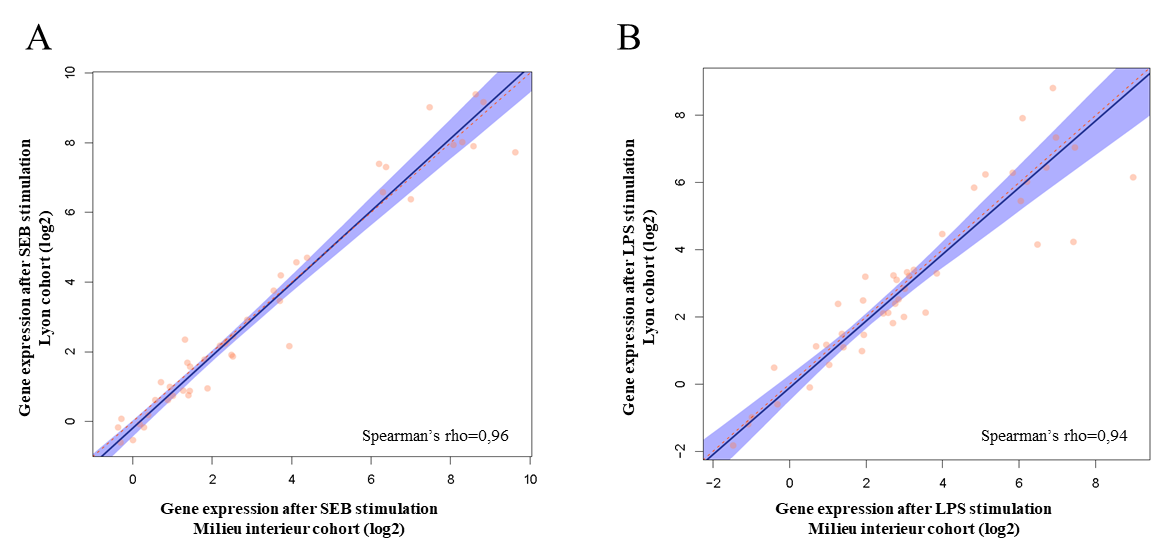

**Figure S1. Comparison between whole-blood response following SEB or LPS stimulation of 20 healthy volunteers from the Lyon study cohort (Y-axis) and from 25 healthy volunteers from the Milieu Intérieur cohort (X-axis).** Comparison of both cohorts was done using a Deming regression method, performed post-SEB (A) or -LPS stimulation (B) on interdecile ranges. A Spearman’s correlation coefficient was calculated for each condition and indicated on each panel. The orange dots represent individual gene expression, the blue beam represents the confidence interval, the blue line the Deming regression and the red dotted line the theoretical 100% identity.

Supplementary table S1

*The 44-gene signature from the Milieu Intérieur study*

| **Gene Name** | **Ensembl gene IDs** | Accession number |
| --- | --- | --- |
| *BST2* | ENSG00000130303 | NM_004335.2 |
| *C3* | ENSG00000125730 | NM_000064.2 |
| *CCL2* | ENSG00000108691 | NM_002982.3 |
| *CCL20* | ENSG00000115009 | NM_004591.1 |
| *CCL4* | ENSG00000129277 | NM_002984.2 |
| *CCL8* | ENSG00000108700 | NM_005623.2 |
| *CCR1* | ENSG00000163823 | NM_001295.2 |
| *CD44* | ENSG00000026508 | NM_001001392.1 |
| *CD83* | ENSG00000112149 | NM_004233.3 |
| *CDKN1A* | ENSG00000124762 | NM_000389.2 |
| *CXCL10* | ENSG00000169245 | NM_001565.1 |
| *CXCL2* | ENSG00000081041 | NM_002089.3 |
| *CXCL9* | ENSG00000138755 | NM_002416.1 |
| *HLA-DMB* | ENSG00000242574; ENSG00000248993 | NM_002118.3 |
| *HLA-DPA1* | ENSG00000231389 | NM_033554.2 |
| *HLA-DPB1* | ENSG00000223865 | NM_002121.4 |
| *HLA-DRA* | ENSG00000204287 | NM_019111.3 |
| *IDO1* | ENSG00000131203 | NM_002164.3 |
| *IFI35* | ENSG00000068079 | NM_005533.3 |
| *IFIH1* | ENSG00000115267 | NM_022168.2 |
| *IFITM1* | ENSG00000185201; ENSG00000185885 | NM_003641.3 |
| *IL1A* | ENSG00000115008 | NM_000575.3 |
| *IL1B* | ENSG00000125538 | NM_000576.2 |
| *IL6* | ENSG00000136244 | NM_000600.1 |
| *IRAK2* | ENSG00000134070 | NM_001570.3 |
| *IRF7* | ENSG00000185507 | NM_001572.3 |
| *JAK2* | ENSG00000096968 | NM_004972.2 |
| *LILRB1* | ENSG00000104972 | NM_001081637.1 |
| *MX1* | ENSG00000157601 | NM_002462.2 |
| *NFKB1* | ENSG00000109320 | NM_003998.2 |
| *NFKB2* | ENSG00000077150 | NM_002502.2 |
| *NFKBIA* | ENSG00000100906 | NM_020529.1 |
| *NFKBIZ* | ENSG00000144802 | NM_001005474.1 |
| *POU2F2* | ENSG00000028277 | NM_002698.2 |
| *RARRES3* | ENSG00000133321 | NM_004585.3 |
| *RELB* | ENSG00000104856 | NM_006509.2 |
| *SLAMF7* | ENSG00000026751 | NM_021181.3 |
| *SOCS1* | ENSG00000185338 | NM_003745.1 |
| *SOCS3* | ENSG00000184557 | NM_003955.3 |
| *SRC* | ENSG00000197122 | NM_005417.3 |
| *STAT2* | ENSG00000170581 | NM_005419.2 |
| *TNFAIP3* | ENSG00000118503 | NM_006290.2 |
| *TNFSF10* | ENSG00000173535 | NM_003810.2 |
| *TNFSF13B* | ENSG00000240505 | NM_006573.4 |
